# CM93, a novel covalent small molecule inhibitor targeting lung cancer with mutant EGFR

**DOI:** 10.1101/2020.03.09.984500

**Authors:** Qiwei Wang, Jing Ni, Tao Jiang, Hwan Geun Choi, Tinghu Zhang, Nathanael Gray, Jean J. Zhao

**Author notes:** Corresponding authors: Jean J. Zhao, Dana-Farber Cancer Institute, 450 Brookline Avenue, Boston, MA 02215.

## Abstract

Epidermal growth factor receptor (EGFR) tyrosine kinase inhibitors (TKIs) have provided successful targeted therapies for patients with EGFR-mutant non-small-cell lung cancer (NSCLC). Osimertinib (AZD9291) is a third-generation irreversible EGFR TKI that has received regulatory approval for overcoming resistance mediated by the EGFR T790M mutation as well as a first-line treatment targeting EGFR activating mutations. However, a significant fraction of patients cannot tolerate the adverse effect associated with AZD9291. In addition, brain metastases are common in patients with NSCLN and remain a major clinical challenge. Here, we report the development of a novel third-generation EGFR TKI, CM93. Compared to AZD9291, CM93 exhibits improved lung cancer targeting and brain penetration and has demonstrated promising antitumor efficacy in mouse models of both EGFR-mutant NSCLC orthotopic and brain metastases. In addition, we find that CM93 confers superior safety benefits in mice. Our results demonstrate that further evaluations of CM93 in clinical studies for patients with EGFR-mutant NSCLC and brain metastases are warranted.

## Introduction

Lung cancer is one of the most common cancers world-wide and a leading cause of cancer deaths. The World Health Organization (WHO) classifies lung cancer into non-small cell lung cancer (NSCLC, 85%) and small-cell lung cancer (SCLC, 15%) with NSCLC comprising squamous, adenocarcinoma and large cell carcinomas^1,2^. Molecular alterations in epidermal growth factor receptor (EGFR) and mutation-caused NSCLC account for over 40% of patients; the remaining 60% include mutations in KRAS, EML-ALK, PTEN, PI3K and others^3,4^.

Epidermal growth factor receptor (EGFR) is a transmembrane protein belonging to the HER/*erb*B family of receptor tyrosine kinases (RTKs), which includes HER1 (EGFR/*erb*B1), HER2 (*neu, erb*B2), HER3 (*erb*B3) and HER4 (*erb*B4)^5^. Hot spot mutations that lead to EGFR overexpression have been associated with a number of cancers including adenocarcinoma of the lung (40% of cases), anal cancers, glioblastoma (50%) and epithelial tumors of the head and neck (80-100%)^6^. EGFR mutations include three classes: Class I mutations include short in-frame deletions that result in the loss of four to six amino acids (E746 to S752) in exon 19; Class II mutations are single-nucleotide substitutions that may occur in exons 18 to 21; Class III mutations are in-frame duplications and/or insertions that occur mostly in exon 20. Among all TK domain mutations, 85–90% are exon 19 class I deletions and exon 21 L858R mutations, which mainly affect sensitivity to EGFR-TKIs^7^.

Gefitinib^8^, Erlotinib^9^ and icotinib^10^ are the first generation EGFR inhibitors approved by the FDA and the Chinese FDA for non-small cell lung cancer (NSCLC). They share a common quinazoline scaffold structure, which reversibly binds to ATP binding sites on over-activated EGFRs to inhibit phosphorylation of EGFR tyrosine residues. Unfortunately, like most other kinase therapies, acquired drug resistance limits their clinical application. Our current understanding of acquired EGFR inhibitor resistance is that most are caused by a secondary mutation converting cytosine (C) at base 2369 is to thymine (T), resulting in the substitution of threonine for methionine at residue 790, but additional mechanisms exist^11,12^. However, the EGFR-T790M mutation accounts for more than 50% of clinical drug-resistant patients. The T790M mutation weakens the ability of Gefitinib/Erlotinib to bind EGFR by altering the spatial conformation of EGFR and increasing the affinity of EGFR to ATP^13^.

The second generation of EGFR inhibitors, which includes Afatinib^14^, Dacomitinib^15^ and Neratinib^16^, are irreversible inhibitors of EGFR and HER2. In addition to competitively occupying ATP binding sites on EGFR, they also alkylate or covalently bind with specific amino acid residues near the opening of the binding pocket of EGFR and thus achieve irreversible inhibition of EGFR. Since the ATP affinity of EGFR T790M is similar to that of WT EGFR, the concentration of quinazoline-based EGFR inhibitors required to inhibit EGFR T790M will also effectively inhibit WT EGFR. In patients this concurrent inhibition of WT EGFR results in skin rash and diarrhea, which limits the dosage that can be utilized to achieve plasma concentrations sufficient to inhibit EGFR T790M^17^.

The 1st covalent pyrimidine EGFR inhibitor targeting the T790M mutation was developed by Dr Pasi A. Jänne and Nathanael Gray at Dana Farber Cancer Institute, with tool compound WZ4002 published in Nature in 2009^18^. This ignited an intense wave of development of a new class of covalent inhibitors attempting to overcome resistance to the first line EGFR treatments, including WZ4002 analogues such as Rociletinib (CO1686)^19^, Olmutinib (BI1482694/HM61713)^20^, Nazartinib (EGF816)^21^ and AZD9291^22^, which was the 1^st^ of the 3rd generation EGFR inhibitors to reach the market and received an ORR of about 70%, 3^rd^ grade AE rate of 17%, drug related death rate of 3.5% (4 cases) in second line therapy after Gerfitinib^23^. WZ4002 class drugs CO1686, and Olmutinib were stopped due to toxicities; EGF816 is still in phase I/II but also facing toxicity issues of skin rashes as well as insufficient efficacy. In 2018, AZD9291 was extended to a 1^st^ line NSCLC trial but one of report disclosed that 34% of patients had a grade 3-4 adverse event; this was still an improvement over Gefitinib at 45%^24^. AZD9291 has been successfully commercialized in the US and some other areas in the world; however, due to its high cost, many NSCLC patients cannot afford this treatment even though it is effective^25^. In addition, approximately 20% of patients cannot tolerate its side effects. Furthermore, the issue of resistance remains due to gatekeeper C797S mutations after 12-18months of continued. Thus, to save the majority of NSCLC patients’ lives, there is a critical need for improved EGFR inhibitors and solutions for the resistances that develop.

We report a new pyrimidine-based molecule termed CM93, a covalent inhibitor, as a third-generation EGFR-TKI for the treatment of EGFR T790M mutation-positive NSCLC. In vitro pharmacology studies revealed that CM93 has good selectivity in kinome and safety panel screening; can efficiently inhibit proliferation of gatekeeper T790M mutation in H1975 cells (EGFR L858R/T790M mutations), and PC9GR4 cells (EGFR del/L858R). In vivo efficacy studies in xenograft mouse tumor models (bearing H1975 or PC9GR4 cells) showed that CM93 has high distribution in the lung and the brain, larger therapeutic window, and an acceptable toxicity profile. These features provide significant benefits for improving therapies against resistance to 1^st^ and 2^nd^ generation EGFR inhibitors. Given these encouraging results, we plan to evaluate CM93 for the next step in development.

## Results

### CM93 is a selective inhibitor against mutant EGFR

To uncover any advantage of CM93 in treating EGFR mutant NSCLC cells, we performed *in vitro* cell proliferation assays using PC9GR4 cells harboring Exon 19 deletion (19Del)/T790M mutations and H1975 cells carrying L858R/T790M mutations and compared results with treatment with AZD9291. Our analysis indicated that CM93 has comparable inhibition efficiency in the PC9GR4 with IC_50_s of 3.66nM for CM93 and 12.03nM for AZD9291; the IC_50S_ for CM93 was 4.39nM vs AZD9291 at 4.8nM in H1975 (Fig. 1A-C).

**Fig 1.**
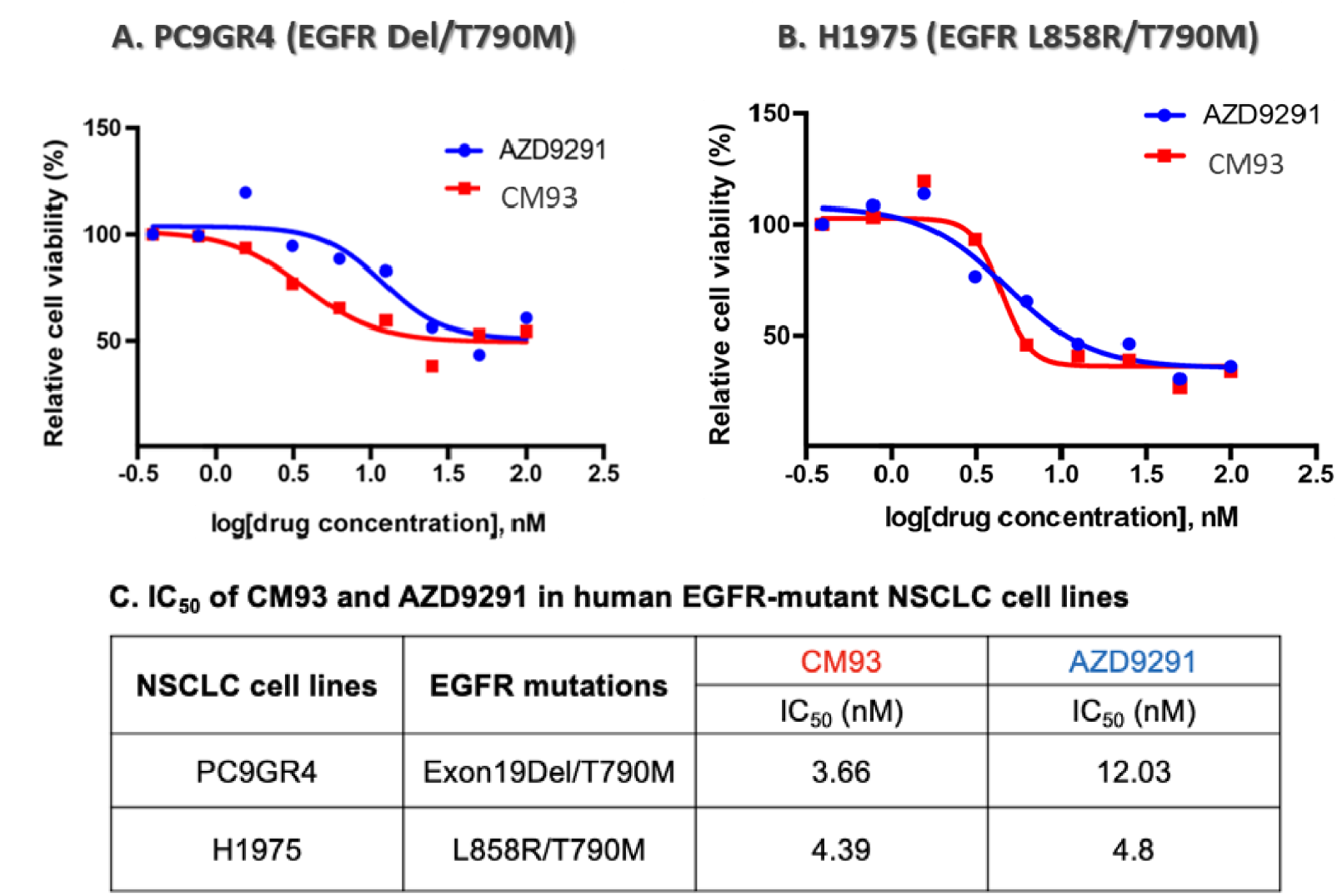
Evaluation of CM93 in EGFR-mutant NSCLC cell lines. (A-C) PC9GR4 (Exon19Del/T790M) and H1975 (L858R/T790M) cells were analyzed by CellTiter Glo following 48 hours of treatment with AZD9291 or CM93.

### CM93 has high tissue distribution in the lung and is effective on regressing tumors in orthotopic mouse models of NSCLC

To confirm *in vivo* inhibition by CM93, we studied CM93 anti-tumor growth inhibition in orthotopic lung cancer using mice bearing H1975 and PC9 cells. Mice were injected via tail vein with H1975-luciferase cells. Beginning 12 days after injection, mice were dosed with CM93 and AZD9291 at 10mg/kg p.o., qd x5 weeks and tumor growth compared with mice in the vehicle control group. Both CM93 and AZD9291 had comparable efficacy and significantly inhibited tumor growth (Fig. 2A). In other experiments we began dosing once large tumors had formed with doses of 25mg/kg for both CM93 and AZD9291. Both drugs efficiently inhibited tumor growth (Fig. 2B).

**Fig. 2.**
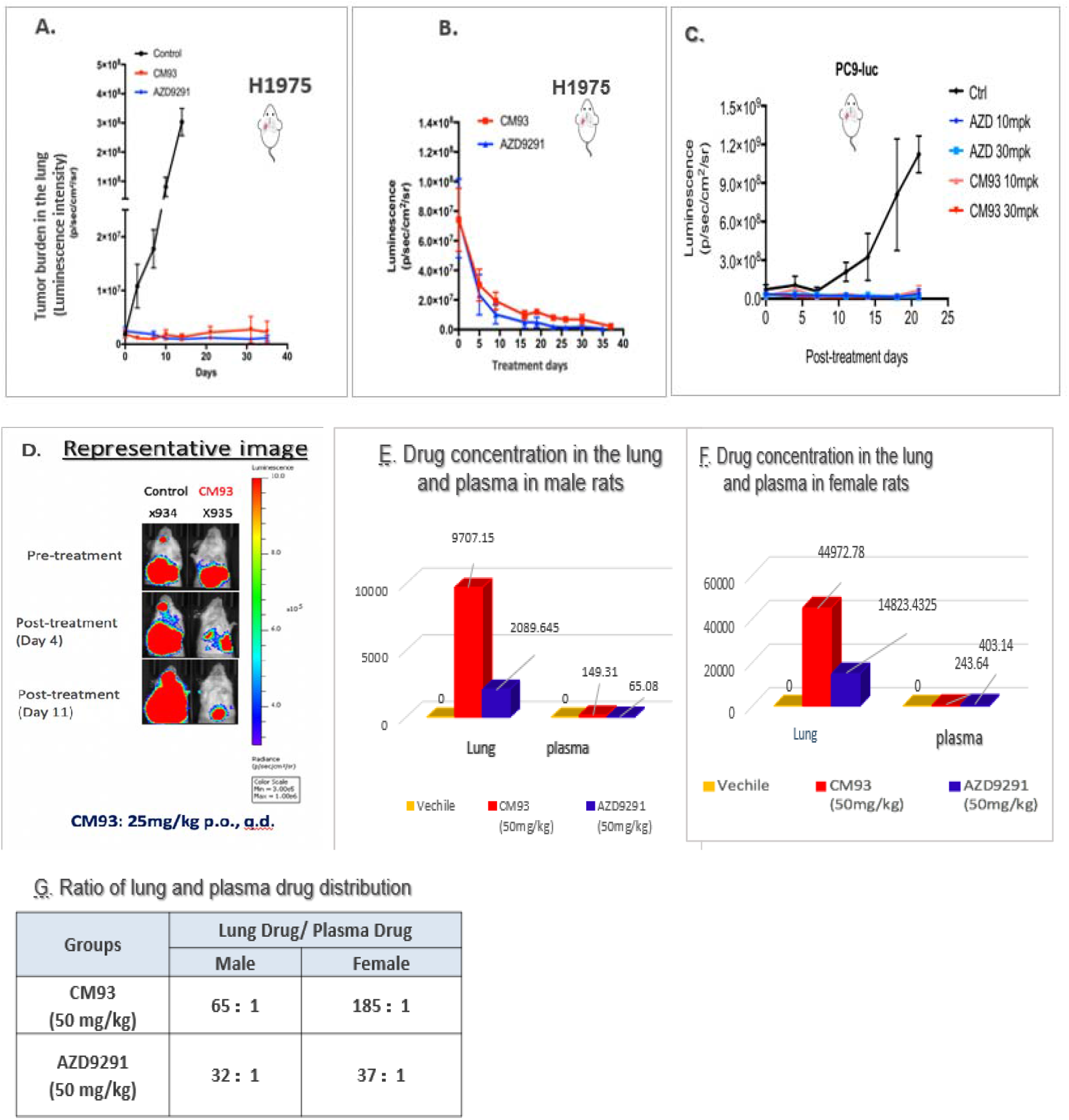
CM93 demonstrates promising antitumor efficacy in orthotopic models of NSCLC. (A-D) Evaluation of CM93 and AZD9291 in orthotopic murine models of NSCLC. 2×10^6^ H1975-luciferase or PC9-luciferase cells were injected into NSG mice via tail vein. (A) 12 days pos-injection, CM93 and AZD9291 were administered at 10mg/kg p.o., qd x5 weeks. (B) Established large tumors were comfirmed by imaging prior to treatment. CM93 and AZD9291 were dosed at 25mg/kg p.o., qd.x 5 weeks. (C) Tail vein injection of PC-9-luciferase cells (2×10^6^), dosing as indicated, po., qd x21days. (D) Tumor imaging results of CM93-treated mice dosed at 25mg/kg for 5 weeks. The tumor (H1975-luc) reduced upon CM93 treatment. (E-G) Lung and plasma distributions of CM93 and AZD9291 (50mg/kg po., qd x 7days) in male (E) and female (F) rats. Drug concentrations were measured by HPLC.

Similar results were seen in a mouse model bearing PC9-luciferase cells; PC-9-luciferase cells (2×10^8^) were injected via tail vein and, after tumor formation (confirmed by imaging), mice were dosed at 10mg/kg and 30mg/kg for both CM93 and AZD9291 groups po., qd x21days. The tumor cell count was significantly reduced at both 10mg/kg and 30mg/kg of CM93 and AZD9291. There was no significant difference between groups treated with CM93 and AZD9291 (Fig. 2C). Fig. 2D was one of the represented imaging of CM93 treated mouse tumor shrinkage followed at pre-treatment, day 4, and day 11’s treatment, in comparison with vehicle mouse.

To further prove the efficacy of CM93 in comparison with AZD9291, we evaluated phosphorylated EGFR levels as a biomarker after dosing of mice subjected to tumor inoculation at different loci using subcutaneous or lung orthotopic inoculation of NSCLC cells (H1975 cells bearing L858/T790M mutations). The results were very striking: phosphorylated EGFR levels were reduced more dramatically by CM93 compared to AZD9291 in the orthotopic NSCLC mouse model, while CM93 reduced phosphorylated EGFR levels less than did AZD9291 in the subcutaneously inoculated model (Data Not Shown). As this indicated differences in tissue distribution of the two drugs, we studied tissue distribution in a rat model by continuous dosing at 50mg/kg for CM93 and AZD9291, qdx7 days (Fig. 2E-G). We then measured drug concentrations in blood and lungs by HPLC. Interestingly, there was a significant difference in the ratio of drug concentrations in lung vs plasma in CM93- and AZD9291-treated animals. CM93 has a much greater lung/plasma ratio than AZD9291: 390:1 vs 193:1 in male rats and 1108:1 vs 220:1 in female rats (Fig. 2E-G). These results demonstrate that both CM93 and AZD9291 are distributed throughout tissues and CM93 is more lung cancer targeted than AZD9291.

### CM93 has high brain penetration and is effective on suppressing the tumor growth in intracranial models of brain metastases

We compared the efficacy of CM93 and AZD9291in an NSCLC brain metastasis model. Mice carrying lung cancer brain metastasis were constructed according to protocol and dosed with CM93 at 25mg/kg and 50mg/kg and AZD9291 at 25mg/kg po qd. x 100 days. CM93 inhibited brain tumor growth in a dose-dependent response pattern compared with both vehicle and AZD9291(Fig. 3A). The 25mg/kg CM93 animals had a medium survival time of 80 days, and 50mg/kg group mice survived over 100days. In contrast, the AZD9291 mice exhibited body weight loss and skin lesions such that they reached an end point after 4 weeks of AZD9291 treatment with a medium survival time of 30 days (Fig. 3B). Both the 25mg/kg and 50mg/kg CM93 treatments were well-tolerated by the mice during the 100 days of dosing while the AZD9291 group mice suffered significant body weight loss when dosed at 25mg/kg (Fig. 3C). These results were supported by brain/plasma drug concentration measurements in rats. After 7 days of continuous dosing with CM93 and AZD9291 at 50mg/kg each, brain and plasma blood sample were prepared for drug concentration assessment by LC-MS/MS analysis. The ratio of brain/plasma drug concentration for CM93 vs AZD9291 was 14:1 vs 6:1 in male rats and15:1 vs 7:1 in female rats (Fig. 3D-F).

**Fig 3.**
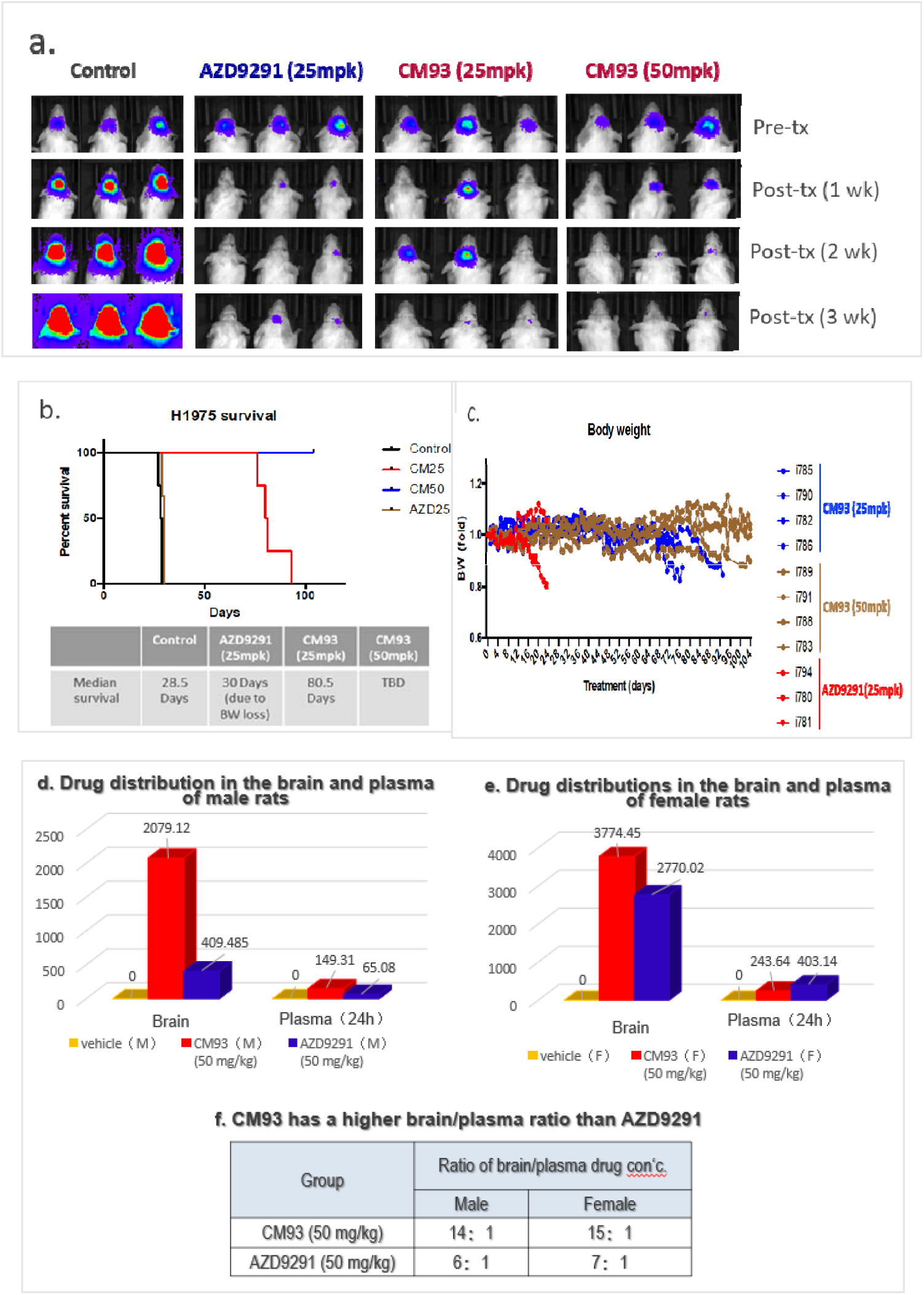
CM93 targets the brain more effectively in a mouse model of H1975 NSCLC brain metastases. Evaluation of CM93 and AZD9291 in SCID mice bearing EGFR-mutant NSCLC brain metastases (H1975-luc). (A) Imaging of CM93 treated mice with EGFR-mutant NSCLC brain metastases in comparison with AZD9291; (B) survival curves of brain metastases mice after treatment of CM93 and AZD9291; (C) Body weight (BW) loss in tumor-bearing mice treated with AZD9291 (25mpk) or CM93 (25 and 50 mpk). (D-F) Brain and plasma distributions of CM93 and AZD9291 (50mg/kg po., qd x 7 days) in male (D) and female (E) rats. Drug concentrations were measured by HPLC.

### CM93 has little adverse effect on mouse skin

Preclinical efficacy studies demonstrated that CM93 is much safer than AZD9291 as assessed by body weight and hair loss as well as development of skin rashes in SCID mice bearing either H1975(T790M) or PC-9(19Del/L858R) tumors and undergoing treatment. We observed that, when treated with AZD9291 at 25mg/kg, qd x30 days, mice lost more than 20% body weight and therefore reached the end point delineated by animal welfare protection requirements. These mice also suffered severe hair loss after three weeks.

In another experiment mice were treated with 10mg/kg (an effective dose for both drugs) CM93 and AZD9291 for 65 days. By the end of the experiment, mice treated with AZD9291 showed more severe weight and hair loss than those treated with CM93. Indeed, mice treated with CM93 showed no hair loss (Fig. 4).

**Fig. 4.**
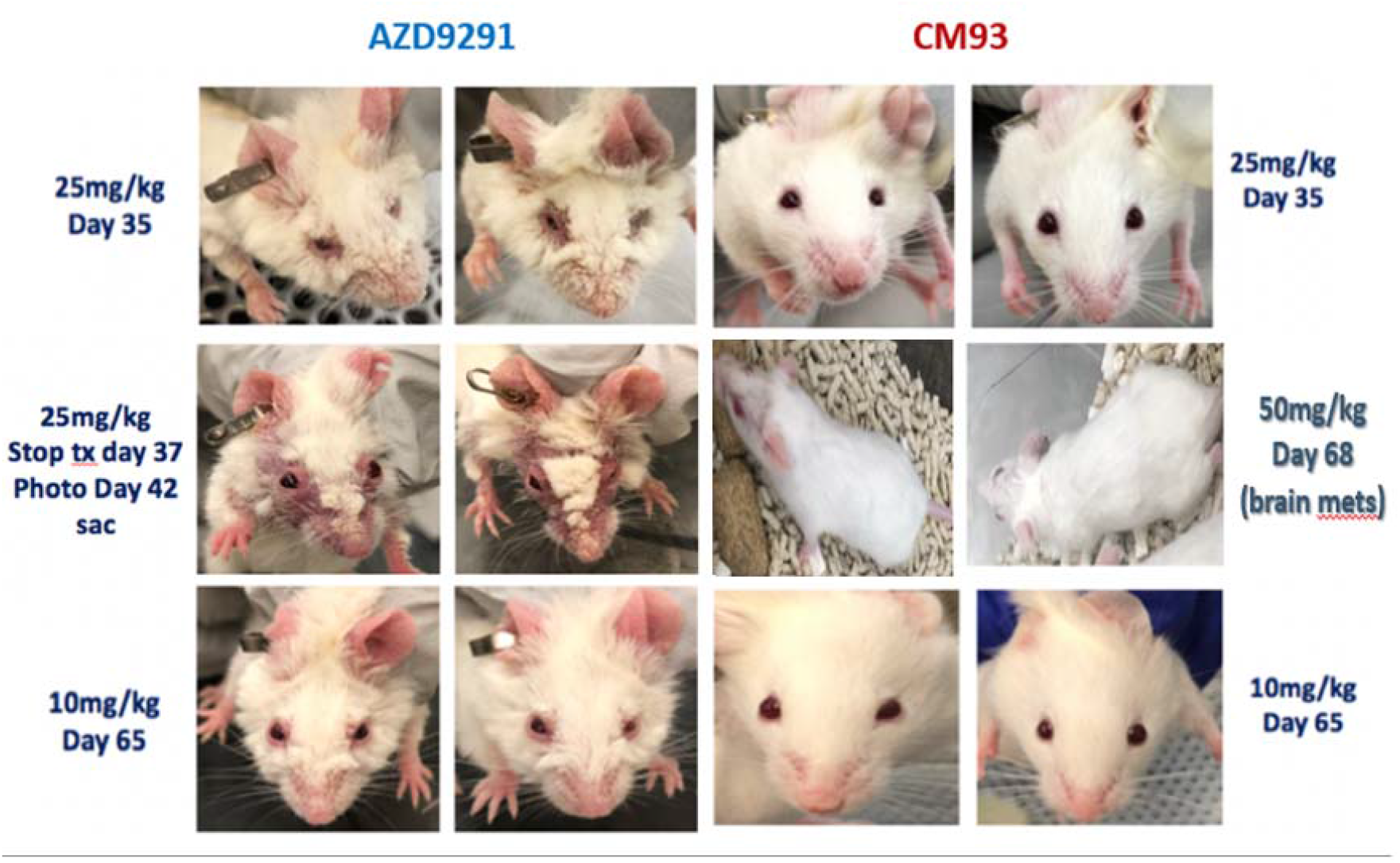
CM93 has little effect on mouse skin. Skin lesions and hair loss were observed in AZD9291-treated but not CM93-treated mice.

## Discussion

Following the 2009^18^ publication of the third generation EGFR inhibitor WZ4002, which contained a pyrimidine-based scaffold, a new round of R&D exploded in the drug industry searching for a compound that could overcome the Exon19Del/T790M mutation. AZD9291 was the one drug among many others that gained extensive commercial success since 2015. However, approximately 20% of patients treated with AZD9291 have intolerable adverse effects, e.g. skin rashes and diarrhea. In recent reports, when AZD9291 was extended to front-line treatment for NSCLC patents, the 3^rd^ and 4^th^ grade adverse event (AE) increased to 34% of patients^24^.

One of the advantages of CM93 is its high selectivity in targeting mutant EGFR in lung cancer. We have compared CM93 to AZD9291 in the treatment of mice bearing xenografts of H1975 cells. Our analysis of the tissue distribution of CM93 in rats showed that CM93 has a higher lung tissue distribution than AZD9291. This higher tissue specific distribution in lung makes CM93 an ideal therapeutic for NSCLC patients.

Another advantage of CM93 in comparison with AZD9291 is brain penetration. Based on our results above, for treatment of NSCLC brain metastasis patients, CM93 is advantageous in that higher doses can be tolerated and it provides a longer median survival rate, which is excellent for future clinical applications.

In summary, we have done a preliminary evaluation of CM93 activity on EGFR mutations *in vitro* and *in vivo* using mice bearing xenograft tumors expressing EGFR T790M and EGFR del/L858R mutations. Our studies have demonstrated the potential advantages of CM93 over the currently-marketed AZD9291 for clinical applications in the future. These advantages include a higher lung and brain tissue selectivity and distribution for CM93 compared to AZD9291, although it has comparable efficacy in tumor growth inhibition especially in a subcutaneous xenograft mouse model. The positive safety profile likely arising from CM93’s lower activity against wild type EGFR will make CM93 more tolerable for patients’ tolerability and improve compliance thus increasing patients’ quality of life. Based on these findings, we plan to move CM93 into further preclinical studies for the future benefit of NSCLC patients.

## Summary of materials and methods

### 1) Kinase inhibitors

Olsimertinib (AZD9291) was obtained from commercial sources. The CM93 compound was synthesized using a four-step chemical synthesis by Pharmaron. The final products were verified by 1H nuclear magnetic resonance and liquid chromatography–mass spectrometry (HPLC/MS).

### 2) Cell lines

*EGFR* wild-type and mutant NSCLC cell lines A549, H1975, PC9 and PC9GR4 were kindly provided by Dr. Pasi A. Jänne at Dana-Farber Cancer Institute (DFCI). All cell lines were authenticated at DFCI and were grown in RPMI-1640 containing 10% FBS (Gibco). Cell viability was determined by CellTier-Glo (Promega). Enzyme kinetic assays were performed *in vitro* using EGFR mutant (L858R/T790M) or WT recombinant proteins and an ATP/NADH coupled assay system as previously described^26^. To investigate the effects of CM93 and AZD9291 on EGFR, AKT and ERK signaling, blood samples harvested from mice following three days’ treatments were analyzed by Western blot.

### 3) Mouse studies

All studies were approved by the Dana-Farber Cancer Institute Animal Care and Use Committee.

To establish the subcutaneous xenograft models of NSCLC, 5×10^6^ H1975 (T790M/L858R) or PC9GR4 (T790M/Exon19del) were subcutaneously impanted in nude mice in 100 microliters serum-free DMEM containing 50% matrigel (Corning)^27^.

In order to monitor tumor growth in the lung or brain by measuring bioluminescence signals in tumor cells, H1975 and PC9 (Exon19del) were transduced with lentiviral vectors expressing the luciferase gene (H1975-luc and PC9-luc). The orthotopic lung tumor models were developed in 8- to 10-week-old NSG mice by tail vein injection of 2×10^6^ cells in 200 microliter PBS per mouse.

For the brain metastases model, H1975-luc or PC9-luc cells (100,000 viable cells in one microliter/mouse) were implanted into 8- to 10-week-old SCID mice via intracranial injections^28^. Mice were treated either with vehicle (0.05N HCL + 0.5% HPMC) alone or CM93 at 1, 3, 10, 30 mg/kg by gavage daily. This was followed by bioluminescent imaging, immunohistochemistry and immunoblotting analyses as previously described. Efficacy was evaluated by % of tumor growth inhibition (%TGI) and % of body weight changes and hair loss and general conditions were documented as indicators of drug toxicity to compared with AZD9291.

For blood, tumor, lung and brain tissue drug concentration analysis, CM93 and AZD9291 were given to rats at 50mg/kg qd for 7days. Samples of blood, tumor, lung and brain tissue were taken at 4 hours after the last dose was given. The concentrations of CM93 and AZD9291 were analyzed on a CTC Autosampler + Agilent 1200+API 4000 LC-MS/MS system.

### 4) Statistical analysis

Statistical significance was determined using unpaired Student’s *t*-tests or ANOVA by GraphPad Prism 6 (GraphPad Software). Data are considered significant when *P* values are < 0.05.

## Disclosure of potential conflicts of interest

Q.W. is a scientific consultant for Crimson Biotech. J.J.Z. is a founder and director of Crimson Biotech and Geode Therapeutics.

## Notes

### Summary of Updates

Figure 1 revised: given that Ba/F3 cells may not represent EGFR-wild type cells, we removed Figure 1D in the original version and edited figure legend and text accordingly.

